# Transient perturbation of the left temporal cortex evokes plasticity-related reconfiguration of the lexical network

**DOI:** 10.1101/562827

**Authors:** Jana Klaus, Dennis J.L.G. Schutter, Vitória Piai

## Abstract

Language impairment is common after left-hemisphere damage. However, the involvement of perilesional and homologous contralateral regions in compensating for left-sided lesions remains poorly understood. The aim of this study was to examine acute organizational changes in brain activity related to conceptual and lexical retrieval in unimpaired language production following transient disruption of the left middle temporal gyrus (MTG). In a randomized single-blind within-subject experiment, we recorded the electroencephalogram from sixteen healthy participants during a context-driven picture-naming task. Prior to the task, the left MTG was perturbed with real neuronavigated continuous theta-burst stimulation (cTBS) or sham stimulation. During the task, participants read lead-in sentences that created a constraining (e.g. “The farmer milks the”) or non-constraining context (e.g. “The farmer buys the”). The last word was shown as a picture that participants had to name (e.g. “cow”). Replicating behavioral studies, participants were overall faster in naming pictures following a constraining relative to a non-constraining context, but this effect did not differ between real and sham cTBS. Real cTBS, however, increased overall error rates compared to sham cTBS. In line with previous studies, we observed a decrease in alpha-beta (8-24 Hz) oscillatory power for constraining relative to non-constraining contexts over left temporal-parietal cortex after participants received sham cTBS. However, following real cTBS, this decrease extended towards left prefrontal regions associated with both domain-general and domain-specific control mechanisms. Our findings provide evidence that immediately after the disruption of the left MTG, the lexical-semantic network is able to quickly reconfigure, also recruiting domain-general regions.

---

Although language function is often severely affected by left-hemisphere brain damage, the mechanisms of cortical reorganization following lesions remain debated. For example, research into recovery from language impairment following a stroke to one or more nodes of the left-hemispheric language network remains equivocal about which reorganization mechanisms are most efficient for language recovery. In particular, it is still debated whether recruitment of homotopic contralateral (i.e. right-hemispheric) areas after left-hemispheric stroke is adaptive or maladaptive for language recovery (Cocquyt, De Ley, Santens, Van Borsel, & De Letter, 2017; Heiss, Kessler, Thiel, Ghaemi, & Karbe, 1999; Meinzer & Breitenstein, 2008; Musso et al., 1999; Naeser et al., 2004, 2005; Rosen et al., 2000; Saur et al., 2006; Thiel et al., 2006; Turkeltaub et al., 2012; Winhuisen et al., 2005, 2007). Additionally, it has been proposed that domain-general systems may help compensate for focal, domain-specific dysfunction (Geranmayeh, Brownsett, & Wise, 2014; Hartwigsen, 2018).Within this framework, lesions in language-relevant regions trigger an upregulation of intact, domain-general networks, particularly within the so-called Multiple Demand Network (MDN). The MDN is generally assumed to be engaged in tasks requiring general cognitive abilities like inhibition, attentional control, cognitive flexibility, and intelligence, necessitating top-down control (Duncan & Owen, 2000; Duncan, 2010; Fedorenko, Duncan, & Kanwisher, 2012), but is not involved in overlearned tasks. Supporting this framework, activity in prefrontal regions has been related to increased task difficulty, both in healthy and brain-lesioned individuals (Geranmayeh et al., 2014; Piai, Roelofs, Acheson, & Takashima, 2013; Vaden et al., 2013). Assuming that lesions to language-specific brain regions likewise increase the difficulty to perform a linguistic task, domain-general regions could very well compensate for (part of) the required functioning to support behavior, minimizing performance impediments.

Importantly, the majority of the empirical work regarding cortical reorganization following lesions has been conducted using functional magnetic resonance imaging, which provides an indirect (i.e., metabolic) measure of neuronal activity (Logothetis & Wandell, 2004) on a rough temporal scale in the order of seconds. As such, it remains largely unknown whether the activity observed outside of the left-hemisphere language network is concurrent with task-related, language-network activity or whether it emerges *after* the language system has failed. Moreover, it is also unclear how these additional areas are integrated as part of a network. These questions can be answered using electrophysiological measures, and in particular oscillations, which provide a window into the dynamic formation of functional networks at the subsecond time scale relevant for language processes. By rhythmically modulating neuronal excitability, neural oscillations coordinate communication between and within distributed neuronal assemblies (Buzsáki & Draguhn, 2004). Using neuronal oscillations thus enables us to investigate the immediate reorganization processes taking place after a focal perturbation.

In the current study, we combined for the first time a perturbation approach to healthy speakers’ brains with electroencephalography (EEG) at the scalp and source levels during a picture-naming task. Prior to the task, real or sham transcranial magnetic stimulation (TMS) was applied to the left middle temporal gyrus (MTG), a key region for lexical retrieval (Baldo, Arévalo, Patterson, & Dronkers, 2013). Real TMS causes a controlled, focal perturbation of the target region, which is not confounded by long-term reorganization in the chronic phase of a language disorder. In combination with EEG, this allows for a time-sensitive neuronal investigation of immediate neuroplastic effects in the otherwise undamaged brain. Note that, although cTBS does not prompt changes in neural states equivalent to those produced by stroke, it does cause lasting suppression of neuronal excitability in the targeted region (Siebner & Rothwell, 2003) of about 50 minutes following application (Wischnewski & Schutter, 2015). Thus, it serves as a proxy to study transient downregulation of a specific area in healthy brain networks.

Behaviorally, we expected a performance decrease in the naming task following real as opposed to sham TMS as a direct marker of the disturbance of the language network. The crucial question was how the oscillatory power modulation in the alpha and beta bands (8-25 Hz), as previously observed in the healthy and reorganized brain after left-hemispheric lesions (Piai, Meyer, Dronkers, & Knight, 2017; Piai, Roelofs, & Maris, 2014; Piai, Roelofs, Rommers, & Maris, 2015; Piai, Rommers, & Knight, 2018), would be affected by the focal perturbation. Following lesion evidence, the oscillatory pattern might shift entirely to the right hemisphere. Using the same task as in the current study, Piai et al. (2017) observed power decreases in the alpha-beta range in the intact right hemisphere in patients with left temporal lesions, suggesting that contralateral regions are recruited when left-hemispheric language nodes are damaged. Alternatively, perilesional networks might get activated in response to the acute perturbation, indicating that acute cortical reorganization is confined to the lesioned hemisphere, where function loss is compensated by other network nodes (Hartwigsen, 2018).

## Methods

### Participants

Following previous studies (Piai et al., 2014, 2015), we recruited 16 participants. We calculated the smallest population effect size we would be able to detect with this sample size at an alpha-level of .05 and 80% power (with the R *pwr* package, Champely 2017), which was *d* = .749. The present study was deemed sufficiently powered since the behavioral effect in previous studies had an effect size of *d* > 2.29 (Piai et al., 2017, 2015) and the EEG effect has a typical effect size of *d* > .80 (Piai et al., 2015).

All participants were right-handed, native Dutch speakers (2 male, mean age = 23.0 years, *SD* = 3.7). Exclusion criteria were a family history of epilepsy, an average use of more than three alcoholic beverages daily, use of psychotropic medication or recreational drugs, skin disease, pregnancy, serious head trauma or brain surgery, neurological or psychiatric disorders, large and/or ferromagnetic metal parts in the head (except for a dental wire), implanted cardiac pacemaker or neurostimulator. All participants gave written informed consent prior to the study, which was approved by the local ethics committee of the Radboud University Medical Centre in Nijmegen (NL64141.091.17).

### Materials

We employed the context-driven picture naming task used in previous studies (Piai et al., 2017, 2014, 2015, 2018). 200 pictures were selected which served as target stimuli. Each picture was associated with two sets of sentences for which the picture names were the last word of the sentences. In the constraining condition, sentences were chosen such that the picture name was highly expected as the final word of the sentence (e.g. “the farmer milks the”), whereas in the non-constraining condition, no one particular word was expected in this position (e.g. “the farmer buys the”). Pictures were allocated to two experimental lists (100 pictures per list corresponding to 200 sentences) to avoid picture repetition across the two experimental sessions. There was no significant difference in sentence length between experimental conditions (constraining: *M* = 6.86, *SD* = 1.87; non-constraining: *M* = 6.73, *SD* = 1.69; *p* > .109) or experimental lists (list 1: *M* = 6.86, *SD* = 1.87; list 2: *M* = 6.73, *SD* = 1.69; *p* > .449).

### Design and Procedure

The design consisted of the two factors sentence context (constraining vs. non-constraining) and stimulation condition (real vs. sham). Sentence context and stimulation condition were fully crossed and tested within participants, with the order counterbalanced between sessions.

Stimulus presentation and response recording was controlled by Presentation (Neurobehavioral Systems, Albany, CA). After EEG preparation, individual resting motor threshold (RMT) for the left hemisphere was determined, followed by the application of neuronavigated cTBS. Immediately afterwards, participants were seated in front of a computer screen. After a short practice block, in which participants were trained to read the sentences and name the pictures without collateral blinking, the experimental task was performed in eight blocks each containing 25 trials. At the beginning of an experimental trial, a fixation cross was presented for 500 ms. Then, each word of the sentence was presented for 300 ms, separated by a blank screen of 200 ms. After the last word, a blank screen appeared for 800 ms, followed by the presentation of the target picture for 1000 ms. Before the next trial was initiated, three asterisks were presented in the center of the screen for 2000 ms, indicating that participants could blink during this period.

#### EEG acquisition

EEG was recorded from 32 Ag/AgCl preamplified scalp electrodes (Biosemi, Amsterdam, The Netherlands) mounted in an elastic cap according to the extended 10-20 system. EEG was sampled at 1,024 Hz. The electrooculogram was recorded horizontally from two electrodes placed on the external canthi of both eyes, and vertically from Fp2 and an electrode placed below the right eye.

#### Continuous theta-burst stimulation

Neuronavigated cTBS (Localite, Sankt Augustin, Germany) was used to navigate the TMS coil and maintain its exact location and orientation for the duration of the stimulation. A figure-of-eight-shaped coil (double 75 mm; coil type MCF-B65) connected to a MagPro X100 stimulator (MagVenture, Farum, Denmark) was used in all cTBS conditions. The stimulation site was based on MNI coordinates corresponding to the region in which all of the patients in Piai et al. (2017) showed damage (i.e., left MTG; MNI: *x* = ‒63, *y* = ‒26, *z* = ‒2). The participants’ position and skull shape was registered in space and transformed to a standard brain, allowing for a precise localization of the target region. Session order (real and sham cTBS) was counterbalanced across participants. During real cTBS, we applied 600 pulses at 50 Hz in trains of three pulses at an inter-burst interval of 200 ms for 40 s. For sham cTBS, the same protocol was administered, but the coil was tilted 90 degrees to mimic the auditory sensation of real cTBS while at the same time preventing current from entering the brain.

Stimulation intensity was set at 80% of the individual RMT of the left hemisphere. Individual RMT was determined using a standardized estimation procedure (Schutter and van Honk 2006). Participants were seated upright and asked to place the arm contralateral to the stimulation site on the upper leg with the palm of the hand facing upwards. The coil was initially placed over M1. By moving the coil in different directions by approximately 1 cm and gradually increasing TMS intensity, the site for eliciting reliable thumb twitches (five out of five) was localized. Next, intensity was decreased until five out of ten consecutive pulses induced a visually identifiable twitch. Finally, the coil was moved again over the scalp and single TMS pulses were applied to make sure no additional scalp site that surpasses the 50% thumb movement criterion was overlooked. If such a site was found, TMS intensity was further decreased according to the 50% criterion. Note that this procedure presumably leads to somewhat higher estimates of individual RMT than measuring RMT on the basis of motor-evoked potentials. However, participants wore the EEG cap during MT estimation and cTBS application, effectively weakening the intensity of the applied stimulation by about 10%. Mean RMT values were 58.96 % (*SD* = 8.16; mean realized coil current gradient = 90 A/us, range: 77 – 103) of mean stimulator output (MSO), corresponding to an average stimulation intensity of 47.25 % (*SD* = 6.56; mean realized coil current gradient = 72 A/us, range: 61 – 83).

#### Behavioral analysis

All analyses were performed using R (version 3.4.1; www.r-project.org). Responses were coded offline for accuracy, and trials in which a wrong or no utterance was produced, or where an utterance was corrected, were removed from the RT analysis. For correct responses, naming latencies were measured manually using Praat (Boersma & Weenink, 2018). Naming latencies were analyzed using linear mixed effects models in the lme4 package (version 1.1.13; (Bates, Mächler, Bolker, & Walker, 2015)). Error rates were analyzed using generalized linear mixed effects models (GLMEM). For all analyses, we included by-participant intercepts to account for interindividual variability in overall task performance, as well as by-participant slopes for the main effect of cTBS. Additionally, we included a by-participant and by-item slope for sentence context. The α-level was set to .05 (two-tailed) for all analyses.

#### EEG analysis

All analyses were performed using FieldTrip version 20171203 (Oostenveld, Fries, Maris, & Schoffelen, 2011) in MatlabR2017a. Trials removed from the RT analysis were also removed from the EEG analysis. Each electrode was re-referenced offline to averaged mastoids. The data were high-pass filtered at 0.16 Hz and segmented into time epochs corresponding to a time window ranging from 1000 ms pre-picture onset to 300 ms post-picture onset. All epochs were inspected individually for eye movements, blinks, and other artefacts, and trials containing artefacts were removed from the analysis (236 trials, 3.7%). Furthermore, excessively noisy channels in individual participants were repaired by spherical spline interpolating (Perrin, Pernier, Bertrand, & Echallier, 1989). Subsequently, time-frequency representations were created and power was calculated with a modified spectrogram approach for frequencies ranging from 2 to 30 Hz at the single-trial level. An adaptive sliding time window of three cycles length was used. Time-frequency representations were then averaged per participant and context by stimulation condition.

#### Source-level analysis

For the source-level analysis, the scalp data were re-referenced offline using the common average reference. Source-level power was estimated for each participant based on the significant scalp-level effect (see below), i.e., −700 ms to −100 ms relative to picture onset and 16 Hz center frequency, using the dynamic imaging of coherent sources method (Gross et al., 2001). A standard boundary element method volume conduction model was used (Oostenveld, Stegeman, Praamstra, & van Oosterom, 2003). The volume was discretized into a grid (1 cm resolution) and the leadfield matrix was calculated for each grid point. The cross-spectral density matrix was computed between 8 and 24 Hz (i.e. spectral smoothing of 8 Hz). The cross-spectral density and leadfield matrices were used to compute common spatial filters (i.e. over both conditions) at each location of the 3D-grid. The common spatial filters were subsequently applied to the Fourier transformed data from each condition separately to obtain source-level spectral power estimates for each grid point. Source-level spectral power estimates were averaged per participant and context by stimulation condition.

#### EEG statistical analyses

The differences in spectral power between conditions for each stimulation type were evaluated using a non-parametric cluster based permutation procedure, which effectively controls the false alarm rate, at both the scalp and source levels (Maris & Oostenveld, 2007). At the scalp level, the tests were performed on all available channels, time points and frequencies. At the source level, the tests were performed over all grid points. Clusters were identified of adjacent data points that exhibited a similar difference between the two conditions across participants based on a two-tailed dependent-samples *t* tests at an α-level of 0.05. Cluster-level statistics were calculated from the summed *t* values within each cluster. Statistical significance was obtained with Monte Carlo stimulations (1,000 random partitions). We assessed the context effect (constraining versus non-constraining) averaged over both cTBS conditions, and also for each cTBS condition separately. At the scalp level, the interaction between context and cTBS condition was assessed by calculating the relative difference between context conditions for each cTBS condition separately, and then comparing those relative differences directly.

## Results

### Left MTG perturbation increases errors in context-driven word retrieval

Figure 2 displays mean naming latencies and error rates broken down by context condition (constraining vs. non-constraining) and cTBS condition (real vs. sham). Following sham cTBS, participants’ mean RTs were 525 ms (*SD* = 215) in the constraining and 704 ms (*SD* = 180) in the non-constraining condition. Following real cTBS, participants’ mean RTs were 521 ms (*SD* = 204) in the constraining and 695 ms (*SD* = 178) in the non-constraining condition. Responses in the constraining condition were reliably faster than in the non-constraining condition (*β* = 90.44, *SE* = 7.20, *t* = 12.56, *p* < .0001, *d* = 0.92), replicating the context effect from previous studies. There was no difference in overall naming latencies as a function of cTBS condition (*β* = −4.35, *SE* = 4.73, *t* = −0.92, *p* = .372, *d* = 0.04). Furthermore, the size of the context effect was comparable following real and sham cTBS (*β* = 0.19, *SE* = 2.32, *t* = 0.08, *p* = .950, *d* = 0.00).

**Figure 1.**
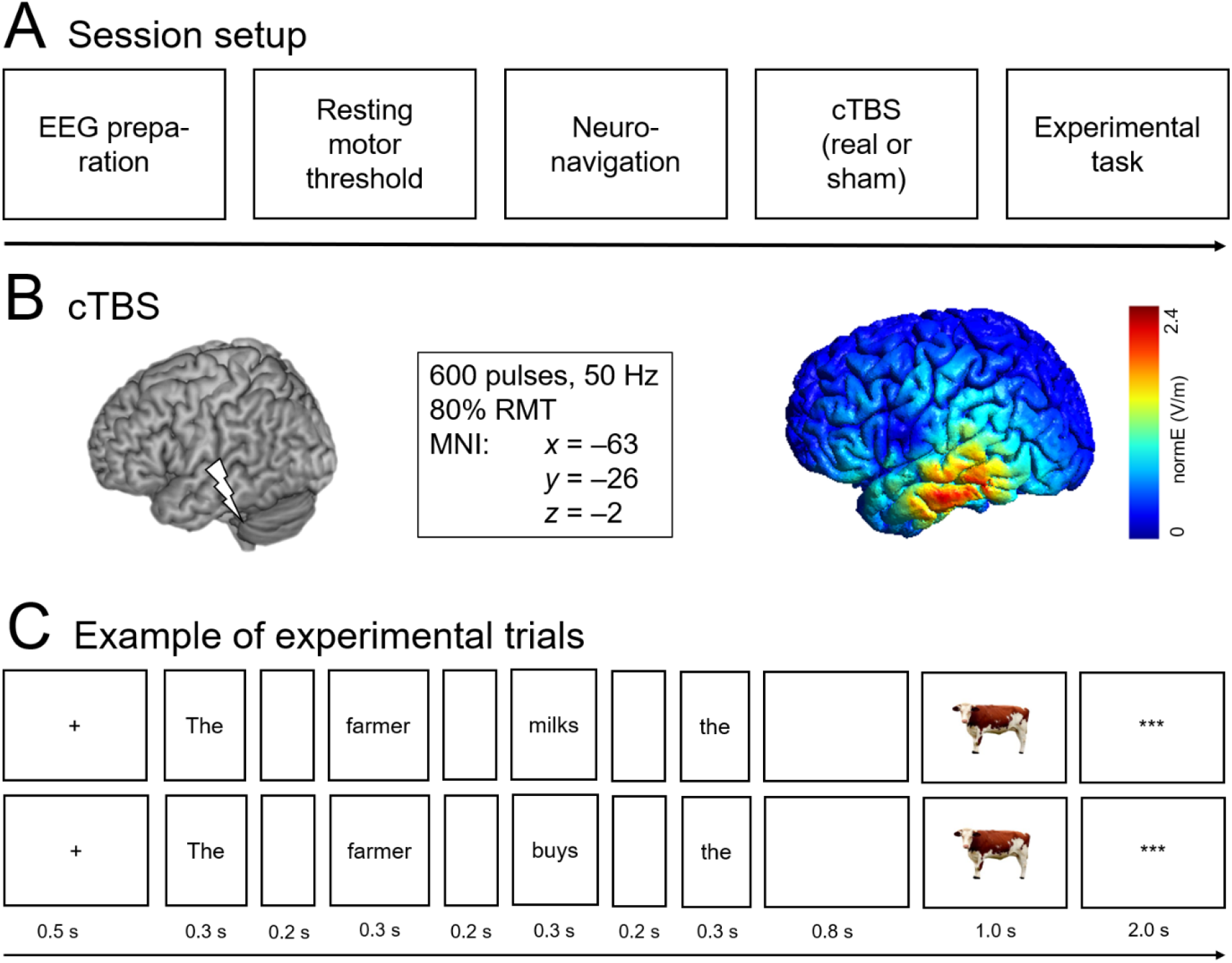
Overview of the experimental procedures. (A) Schematic illustration of an experimental session. (B) Illustration of the cTBS target site (left) and the induced magnetic field as simulated in an example brain using SimNIBS (Thielscher et al. 2015) (right). (C) Two example trials illustrating the constraining context condition (top) and non-constraining context condition (bottom), respectively.

**Figure 2.**
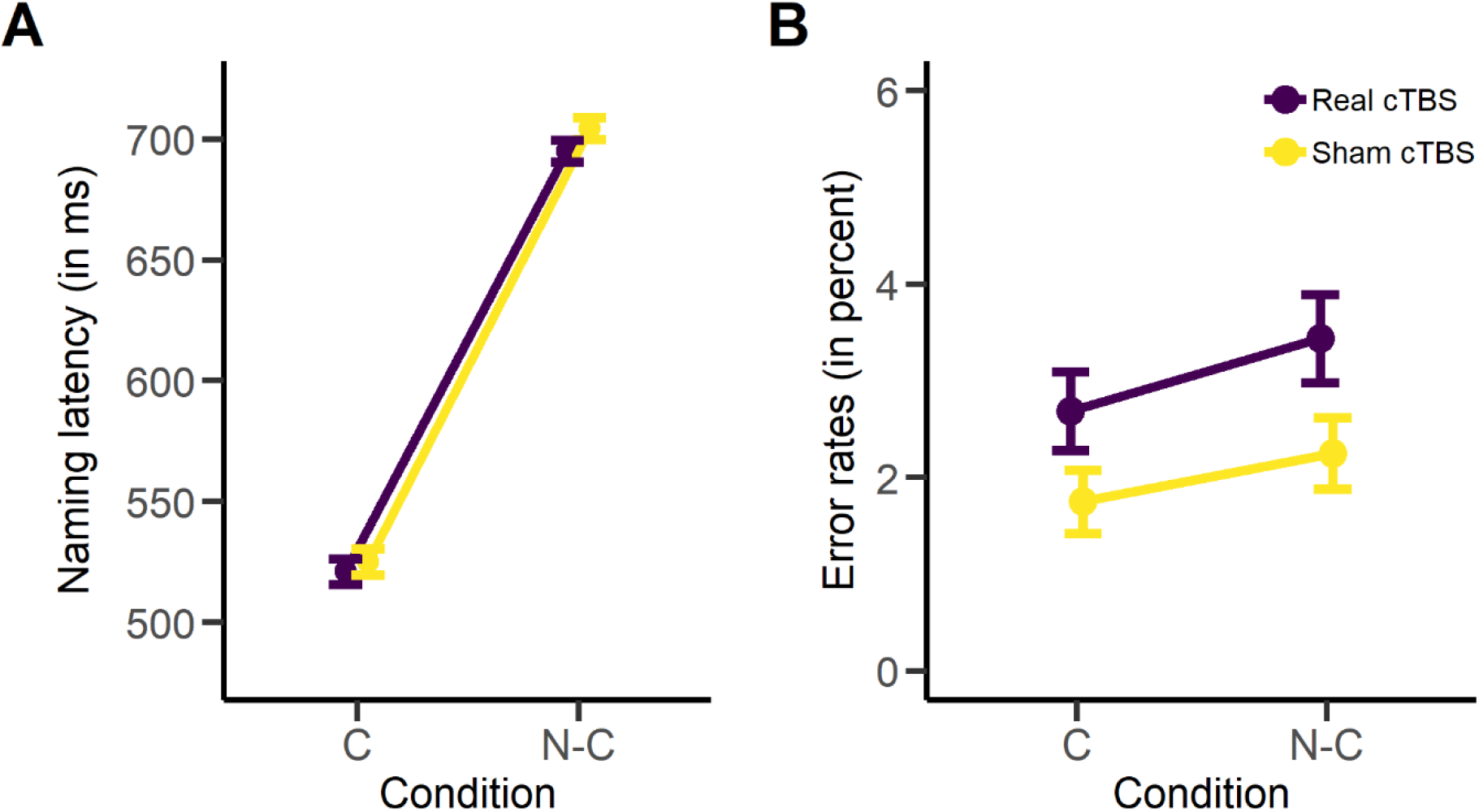
Behavioral results. (A) Mean naming latencies (in ms), broken down by cTBS condition (real vs. sham) and context condition (constraining vs. non-constraining). (B) Error rates (in percent), broken down by cTBS condition (real vs. sham) and context condition (constraining vs. non-constraining). C = constraining; N-C = non-constraining. Error bars represent standard errors of the mean.

Following sham cTBS, participants’ error rates were 1.8% (*SD* = 1.4) and 2.3% (*SD* = 1.5) for the constraining and non-constraining condition, respectively. Following real cTBS, participants made 2.7% (*SD* = 1.7) errors in the constraining and 3.4% (*SD* = 1.8) errors in the non-constraining condition. In the GLMEM analysis, the maximal model as specified in the Methods section did not converge. Thus, we reduced the random-effects structure, which resulted in a final model that contained by-participant and by-item intercepts as well as a by-item slope for sentence context. In this model, error rates did not differ between the constraining and the non-constraining condition (*β* = 0.13, *SE* = 0.08, *z* = 1.60, *p* = .110, odds ratio = 1.14, 95% CI: 0.97 – 1.35). Participants made more errors following real compared to sham cTBS (*β* = 0.23, *SE* = 0.08, *z* = 2.73, *p* = .006, odds ratio = 1.26, 95% CI: 1.07 – 1.49). Context condition and stimulation condition did not interact (*β* = 0.00, *SE* = 0.08, *z* = 0.01, *p* = .995, odds ratio = 1.00, 95% CI: 0.85 – 1.18).

### Left MTG perturbation modulates pre-picture alpha-beta oscillations

Across both real and sham cTBS conditions, power decreases were stronger following constraining relative to non-constraining contexts (Monte Carlo *p* < .004). Moreover, we also assessed the context effect for each cTBS condition separately, as shown in Figure 3. Following sham cTBS, a statistically significant cluster was found (Monte Carlo *p* = .006). Here, power decreases were most prominent between 8 and 24 Hz and between 700 to 100 ms prior to picture onset and extended over left posterior and left and right anterior channels (see left panel of Figure 3), replicating previous findings. By contrast, following real cTBS, this context effect was attenuated, resulting in no significant clusters over the scalp (Monte Carlo *p* = .080). Looking at the group-level time-frequency representations broken down by cTBS condition (Figure 3) reveals differences in the topographical distribution of the context effect, implying a different neuronal configuration of the underlying sources for sham vs. real cTBS. The interaction between cTBS and context condition was not significant (Monte Carlo *p* > .976). However, we note that cluster-based permutation testing is appropriate for assessing the hypothesis of exchangeability across the conditions tested, but not for inferring specific spatial-spectro-temporal differences between conditions (Maris & Oostenveld, 2007). Therefore, to follow up on the spatial differences between cTBS conditions, we localized the neuronal sources of the context effect for each condition separately.

**Figure 3.**
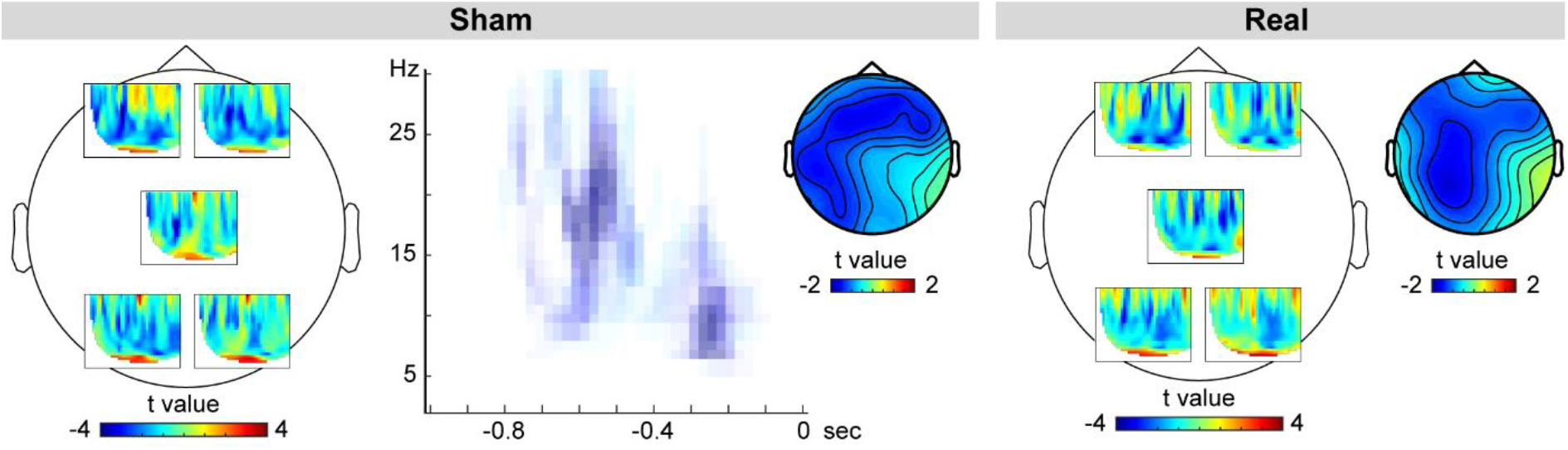
Group-level time-frequency representation and scalp distribution of the power changes for the constraining relative to the non-constraining condition, broken down by cTBS condition. T-values for the comparison between time-resolved spectra are shown for five channels. The location of each time-resolved spectra corresponds roughly to the location of the respective channel. For sham cTBS, the significant cluster is shown averaged over the channels belonging to the cluster; data points not pertaining to the cluster are masked. Scalp topographies are shown for the averages between 8 and 24 Hz and −700 and −100 ms.

### Left MTG perturbation causes additional recruitment of left prefrontal regions

The difference in scalp topographies between the cTBS conditions implies distinct patterns of neuronal generators. To allow for an anatomically more defined comparison between the two cTBS conditions, we source-localized participants’ context effects using frequency-domain beamformers across the time-frequency window found to elicit the strongest power decrease at the scalp level for sham stimulation and for both cTBS conditions combined (i.e. 700 to 100 ms prior to picture onset and from 8 to 24 Hz). Power decreases in this time-frequency range were statistically significant following sham cTBS (*p* = .008) as well as real cTBS (*p* = .002). The source results, displayed as relative power decreases separated by cTBS condition, are shown in Figure 4A. Following sham cTBS, the context effect was localized in left temporal and parietal regions, replicating previous findings from MEG (Piai et al., 2015). By contrast, following real cTBS, this effect was much more widespread towards left prefrontal regions, additionally encompassing the left frontal cortex and anterior temporal lobe. Figure 4B displays the source differences between the two cTBS conditions, illustrating which regions were selectively recruited following sham (blue) and real (red) cTBS. Left prefrontal regions were selectively recruited following real cTBS. By contrast, following sham cTBS, only comparably small portions of the left precentral gyrus were selectively recruited.

**Figure 4.**
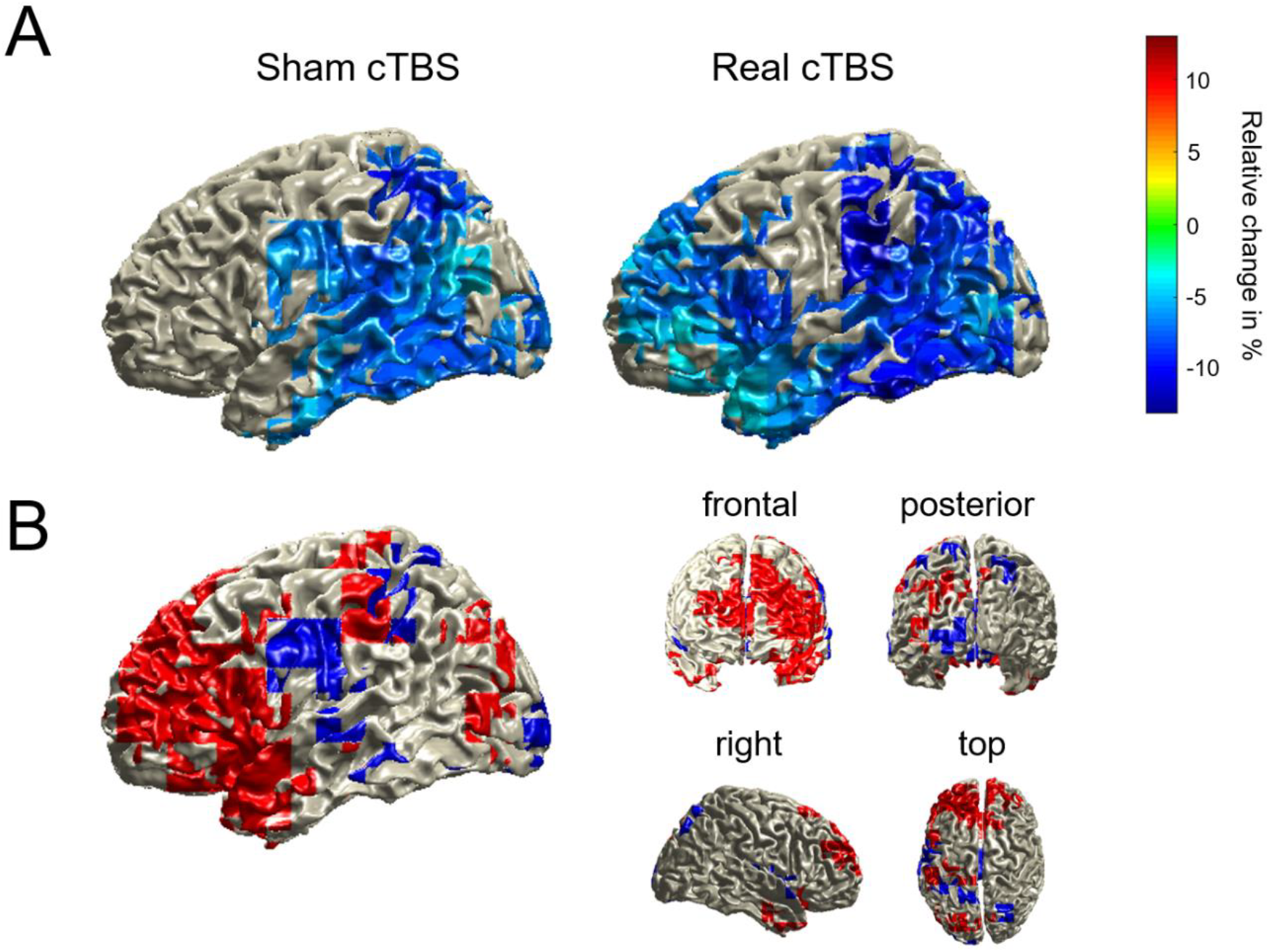
(A) Source localization of the power decreases for constraining relative to non-constraining contexts (displayed in relative percent change) for both cTBS conditions, masked by the statistically significant grid points. (B) Difference plot displaying supra-threshold source-level activity specific for real cTBS (red) and sham cTBS (blue).

## General Discussion

The goal of the present study was to investigate the immediate effects of a downregulation of the mid-section of the left temporal gyrus on the behavioral performance as well as on the time-resolved oscillatory patterns of context-guided language production. Transiently perturbing activity in the left MTG with high-frequency cTBS prior to the task effectively disrupted the contribution of this node within the semantic language production network. Behaviorally, this translated into an increase in overall task difficulty, as shown by an increase in picture naming error rates, regardless of whether the to-be-named picture was expected or not. Picture naming latencies showed the facilitation effect from sentence context irrespective of cTBS condition. Thus, real cTBS affected language *production*, rather than comprehension.

At the neuronal level, the sham condition replicated previous findings of power decreases in the alpha-beta band in left temporal and inferior parietal regions prior to picture onset (Piai et al., 2017, 2014, 2015, 2018). By contrast, the left MTG perturbation resulted in attenuated scalp effects and, at the source level, additional recruitment of left prefrontal regions concurrent with the left posterior activity.

Previous studies have shown that a rise in task difficulty is associated with stronger activity in left ventro- and dorsolateral prefrontal cortex and the cingulo-opercular network, which encompasses the dorsal anterior cingulate cortex, superior frontal gyrus, and sometimes posterior inferior frontal gyrus (Fedorenko, Behr, & Kanwisher, 2011; Fedorenko, Duncan, & Kanwisher, 2013; Fridriksson & Morrow, 2005; Geranmayeh et al., 2014). Our source-level analysis of task-specific oscillatory activity is in line with these previous findings, suggesting recruitment of the domain-general control network in the face of higher task demands. Importantly, using electrophysiological measures and, in particular, analyses of oscillatory activity, we were able to demonstrate that these left prefrontal areas are engaged in the same time scale (i.e. in less than a second) as the left posterior areas, rather than (seconds) later in time. This finding is particularly novel and has substantial implications for understanding mechanisms of network reconfiguration, in that it provides temporally and spatially specific evidence of the concurrent recruitment of ‘perilesional’ regions following a focal perturbation. Moreover, to the extent that oscillatory activity provides a window into the dynamic formation of functional networks (Buzsáki & Draguhn, 2004), the prefrontal regions appear to be integrated into the oscillatory network, facilitating communication across its different nodes by rhythmically modulating excitability of their neurons. It is possible that the vicinity of the stimulated region to the inferior fronto-occipital and inferior longitudinal fasciculi connecting left posterior to anterior regions (Duffay et al., 2005; Turken & Dronkers, 2011) facilitated the spread of activity to frontal regions in the real cTBS condition. In other words, these connections may have mediated the integration of the frontal regions as observed in the oscillatory activity, increasing the size of the network in the same frequency band.

Interestingly, we found no evidence that the right hemisphere performs a compensating role when part of the left-hemispheric network is disturbed, as has been reported for chronic aphasic patients with lesions in the left hemisphere (Musso et al., 1999; Piai et al., 2017; Saur et al., 2006; Turkeltaub et al., 2012; Winhuisen et al., 2005). This suggests that acute responses to focal left-hemispheric damage, which in the current study was imitated by the application of cTBS, were offset entirely by perilesional regions in the same hemisphere as opposed to homotopic regions. Indeed, most studies reporting right-hemispheric compensation patterns are confined to chronic patients, who might have already undergone extensive, more widespread reorganization of language networks. Of course, investigating the immediate (i.e. acute) effect of a lesion in real patients is highly challenging for many organizational and pragmatic reasons. In that sense, study procedures like the current one can provide insights that help determine plausible language-related neuroplasticity mechanisms.

A similar pattern of additional prefrontal cortex recruitment with the same paradigm was observed in an MEG study by Piai et al. (2015), in which healthy volunteers alternated between naming the picture presented at the end of a sentence (i.e. language production, identical to the current task) and judging, via button press, whether the picture was expected or not. This additional task demand (i.e. switching between two different output modes) was reflected in a similar pattern of additional source activity in the prefrontal cortex during the language production task as obtained in the current study. By contrast, Roos and Piai (in preparation) localized alpha-beta oscillatory activity exclusively to the left temporal lobe when participants only executed the naming task, that is, in the absence of additional higher-order control, and comparable to the sham condition in the current study. Combined, these findings provide further evidence that increased left prefrontal activity may be associated with increased executive control involvement.

Arguably, the quality of the source localization in the present study is not optimal, as we did not have exact electrode locations nor individual scans for the reconstruction of the sources (Akalin Acar & Makeig, 2013). Despite this caveat, we are confident in the accuracy of our results given the similarity of the observed pattern in left posterior cortex in the sham condition with previous studies [Piai et al., 2015; Roos and Piai, in preparation]. Moreover, localization errors resulting from these shortcomings should be in the margin of 10-15 mm (Akalin Acar & Makeig, 2013), whereas the anterior spread we observed extends beyond that. Regardless, caution should be observed when interpreting the source locations strictly. Ideally, the results should be replicated in future studies with more carefully computed forward models. Nonetheless, the current results support the inference that an increase in task demands, as caused by perturbing an important node in the lexical-semantic network, triggers an immediate recruitment of additional left prefrontal regions, possibly associated with the Multiple Demand Network.

The discrepancy between the scalp- and source-level results for the real cTBS condition deserves some attention. Using Monte Carlo simulations, no reliable oscillatory effect was found at the scalp level following real cTBS, indicating that the effect became more variable across participants in the spatial-spectro-temporal domain. At the source level, however, the effect was reliable across participants (see also Piai et al. 2015 for a similar apparent discrepancy between scalp- and source-level results). This discrepancy can be explained by the fact that beamforming, the source localization method we employed, is a spatial filtering technique which suppresses the contributions of noise sources that are temporally correlated to reconstruct the neuronal sources of an effect of interest (Gross et al., 2001). The scalp-level activity is a conglomerate of many sources of signal/noise. An improved signal-to-noise ratio is expected once spatial filtering is employed, leading to significant effects at the source level as in the current study.

In summary, the current study showed for the first time how behavioral and underlying electrophysiological fingerprints of language production are affected by transient perturbation of a key region within the semantic network. The findings show that this results in an initiation of adaptive processes of functional networks to left frontal and temporal regions associated with both domain-general as well as domain-specific, lexical-semantic processes. However, this compensation mechanism was not successful in entirely alleviating performance decrements, as overall error rates were moderately, but significantly increased following real cTBS. Together, these results provide new insights into the cortical mechanisms at play in response to disruption of normal functioning.

## Acknowledgments

This work was supported by the German Research Foundation (grant number KL 2933/2-1 to J. K.) and the Netherlands Organization for Scientific Research (grant number 451-17-003 to V. P.). We would like to thank Anna Dewenter for invaluable assistance in data collection and Gesa Hartwigsen for helpful comments on previous versions of this manuscript.

## References

Akalin Acar, Z., & Makeig, S. (2013). Effects of forward model errors on EEG source localization. Brain Topography, 26, 378–396. https://doi.org/10.1007/s10548-012-0274-6

Baldo, J. V, Arévalo, A., Patterson, J. P., & Dronkers, N. F. (2013). Grey and white matter correlates of picture naming: evidence from a voxel-based lesion analysis of the Boston Naming Test. bCortex, 49(3), 658–667. https://doi.org/10.1016/j.cortex.2012.03.001

Bates, D., Mächler, M., Bolker, B., & Walker, S. (2015). Fitting linear mixed-effects models using lme4. Journal of Statistical Software, 67(1), 1–48. https://doi.org/10.18637/jss.v067.i01

Boersma, P., & Weenink, D. (2018). Praat: Doing phonetics by computer. Retrieved from http://www.praat.org/

Buzsáki, G., & Draguhn, A. (2004). Neuronal oscillations in cortical networks. Science, 304(5679), 1926–1929. https://doi.org/10.1126/science.1099745

Champely, S. (2017). pwr: Basic functions for power analysis. R package version 1.2-1. https://CRAN.R-project.org/package=pwr.

Cocquyt, E.-M., De Ley, L., Santens, P., Van Borsel, J., & De Letter, M. (2017). The role of the right hemisphere in the recovery of stroke-related aphasia: A systematic review. Journal of Neurolinguistics, 44, 68–90. https://doi.org/10.1016/j.jneuroling.2017.03.004

Duffay, H., Gatignol, P., Mandonnet, E., Peruzzi, P., Tzourio-Mazoyer, N., & Capelle, L. (2005). New insights into the anatomo-functional connectivity of the semantic system: a study using cortico-subcortical electrostimulation. Brain, 128, 797–810. https://doi.org/10.1093/brain/awh423

Duncan, J. (2010). The multiple-demand (MD) system of the primate brain: mental programs for intelligent behaviour. Trends in Cognitive Sciences, 14, 172–179. https://doi.org/10.1016/j.tics.2010.01.004

Duncan, J., & Owen, A. M. (2000). Common regions of the human frontal lobe recruited by diverse cognitive demands. Trends in Neurosciences, 23(10), 475–483. https://doi.org/10.1016/S0166-2236(00)01633-7

Fedorenko, E., Behr, M. K., & Kanwisher, N. (2011). Functional specificity for high-level linguistic processing in the human brain. Proceedings of the National Academy of Sciences of the United States of America, 108(39), 16428–33. https://doi.org/10.1073/pnas.1112937108

Fedorenko, E., Duncan, J., & Kanwisher, N. (2012). Language-selective and domain-general regions lie side by side within Broca’s area. Current Biology, 22(21), 2059–62. https://doi.org/10.1016/j.cub.2012.09.011

Fedorenko, E., Duncan, J., & Kanwisher, N. (2013). Broad domain generality in focal regions of frontal and parietal cortex. Proceedings of the National Academy of Sciences, 110(41), 16616–21. https://doi.org/10.1073/pnas.1315235110

Fridriksson, J., & Morrow, L. (2005). Cortical activation and language task difficulty in aphasia. Aphasiology, 19(3–5), 239–250. https://doi.org/10.1080/02687030444000714

Geranmayeh, F., Brownsett, S. L. E., & Wise, R. J. S. (2014). Task-induced brain activity in aphasic stroke patients: what is driving recovery? Brain, 137(10), 2632–2648. https://doi.org/10.1093/brain/awu163

Gross, J., Kujala, J., Hämäläinen, M., Timmermann, L., Schnitzler, A., & Salmelin, R. (2001). Dynamic imaging of coherent sources: Studying neural interactions in the human brain. PNAS (Vol. 98).

Hartwigsen, G. (2018). Flexible redistribution in cognitive networks. Trends in Cognitive Sciences, 22(8), 687–698. https://doi.org/10.1016/j.tics.2018.05.008

Heiss, W.-D., Kessler, J., Thiel, A., Ghaemi, M., & Karbe, H. (1999). Differential capacity of left and right hemispheric areas for compensation of poststroke aphasia. Annals of Neurology, 45(4), 430–438. https://doi.org/10.1002/1531-8249(199904)45:4<430::AID-ANA3>3.0.CO;2-P

Logothetis, N. K., & Wandell, B. A. (2004). Interpreting the BOLD signal. Annual Review of Physiology, 66(1), 735–769. https://doi.org/10.1146/annurev.physiol.66.082602.092845

Maris, E., & Oostenveld, R. (2007). Nonparametric statistical testing of EEG- and MEG-data. Journal of Neuroscience Methods, 164(1), 177–190. https://doi.org/10.1016/J.JNEUMETH.2007.03.024

Meinzer, M., & Breitenstein, C. (2008). Functional imaging studies of treatment‐induced recovery in chronic aphasia. Aphasiology, 22(12), 1251–1268. https://doi.org/10.1080/02687030802367998

Musso, M., Weiller, C., Kiebel, S., Müller, S. P., Bülau, P., & Rijntjes, M. (1999). Training-induced brain plasticity in aphasia. Brain, 122, 1781–1790.

Naeser, M. A., Martin, P. I., Baker, E. H., Hodge, S. M., Sczerzenie, S. E., Nicholas, M., … Yurgelun-Todd, D. (2004). Overt propositional speech in chronic nonfluent aphasia studied with the dynamic susceptibility contrast fMRI method. NeuroImage, 22(1), 29–41. https://doi.org/10.1016/J.NEUROIMAGE.2003.11.016

Naeser, M. A., Martin, P. I., Nicholas, M., Baker, E. H., Seekins, H., Kobayashi, M., … Pascual-Leone, A. (2005). Improved picture naming in chronic aphasia after TMS to part of right Broca’s area: An open-protocol study. Brain and Language, 93, 95–105. https://doi.org/10.1016/j.bandl.2004.08.004

Oostenveld, R., Fries, P., Maris, E., & Schoffelen, J.-M. (2011). FieldTrip: Open source software for advanced analysis of MEG, EEG, and invasive electrophysiological data. Computational Intelligence and Neuroscience, 2011, 156869. https://doi.org/10.1155/2011/156869

Oostenveld, R., Stegeman, D. F., Praamstra, P., & van Oosterom, A. (2003). Brain symmetry and topographic analysis of lateralized event-related potentials. Clinical Neurophysiology, 114(7), 1194–1202. https://doi.org/10.1016/S1388-2457(03)00059-2

Perrin, F., Pernier, J., Bertrand, O., & Echallier, J. F. (1989). Spherical splines for scalp potential and current density mapping. Electroencephalography and Clinical Neurophysiology, 72, 184–187.

Piai, V., Meyer, L., Dronkers, N. F., & Knight, R. T. (2017). Neuroplasticity of language in left-hemisphere stroke: Evidence linking subsecond electrophysiology and structural connections. Human Brain Mapping, 38(6), 3151–3162. https://doi.org/10.1002/hbm.23581

Piai, V., Roelofs, A., Acheson, D. J., & Takashima, A. (2013). Attention for speaking: domain-general control from the anterior cingulate cortex in spoken word production. Frontiers in Human Neuroscience, 7, 832. https://doi.org/10.3389/fnhum.2013.00832

Piai, V., Roelofs, A., & Maris, E. (2014). Oscillatory brain responses in spoken word production reflect lexical frequency and sentential constraint. Neuropsychologia, 53, 146–156. https://doi.org/10.1016/j.neuropsychologia.2013.11.014

Piai, V., Roelofs, A., Rommers, J., & Maris, E. (2015). Beta oscillations reflect memory and motor aspects of spoken word production. Human Brain Mapping, 36(7), 2767–2780. https://doi.org/10.1002/hbm.22806

Piai, V., Rommers, J., & Knight, R. T. (2018). Lesion evidence for a critical role of left posterior but not frontal areas in alpha-beta power decreases during context-driven word production. European Journal of Neuroscience, 48(7), 2622–2629. https://doi.org/10.1111/ejn.13695

Roos, N, & Piai, V. Across-session consistency of context-driven language production: a magnetoencephalography study. In preparation.

Rosen, H. J., Petersen, S. E., Linenweber, M. R., Snyder, A. Z., White, D. A., Chapman, M. A., … Corbetta, M. (2000). Neural correlates of recovery from aphasia after damage to left inferior frontal cortex. Neurology, 55, 1883–1894.

Saur, D., Lange, R., Baumgaertner, A., Schraknepper, V., Willmes, K., Rijntjes, M., & Weiller, C. (2006). Dynamics of language reorganization after stroke. Brain, 129(6), 1371–1384. https://doi.org/10.1093/brain/awl090

Siebner, H., & Rothwell, J. (2003). Transcranial magnetic stimulation: new insights into representational cortical plasticity. Experimental Brain Research, 148(1), 1–16. https://doi.org/10.1007/s00221-002-1234-2

Thiel, A., Habedank, B., Herholz, K., Kessler, J., Winhuisen, L., Haupt, W. F., & Heiss, W.-D. (2006). From the left to the right: How the brain compensates progressive loss of language function. Brain and Language, 98, 57–65. https://doi.org/10.1016/j.bandl.2006.01.007

Turkeltaub, P. E., Branch Coslett, H., Thomas, A. L., Faseyitan, O., Benson, J., Norise, C., & Hamilton, R. H. (2012). The right hemisphere is not unitary in its role in aphasia recovery. Cortex, 48, 1179–1186. https://doi.org/10.1016/j.cortex.2011.06.010

Turken, A. U., & Dronkers, N. F. (2011). The neural architecture of the language comprehension network: converging evidence from lesion and connectivity analyses. Frontiers in Systems Neuroscience, 5, 1. https://doi.org/10.3389/fnsys.2011.00001

Vaden, K. I., Kuchinsky, S. E., Cute, S. L., Ahlstrom, J. B., Dubno, J. R., & Eckert, M. A. (2013). The cingulo-opercular network provides word-recognition benefit. Journal of Neuroscience, 33(48), 18979–18986. https://doi.org/10.1523/JNEUROSCI.1417-13.2013

Winhuisen, L., Thiel, Alexander, Schumacher, B., Kessler, J., Rudolf, J., Haupt, W. F., & Heiss, W. D. (2007). The right inferior frontal gyrus and poststroke aphasia: A follow-up investigation. Stroke, 38, 1286–1292. https://doi.org/10.1161/01.STR.0000259632.04324.6c

Winhuisen, L., Thiel, A., Schumacher, B., Kessler, J., Rudolf, J., Haupt, W. F., & Heiss, W. D. (2005). Role of the contralateral inferior frontal gyrus in recovery of language function in poststroke aphasia: A combined repetitive transcranial magnetic stimulation and positron emission tomography study. Stroke, 36, 1759–1763. https://doi.org/10.1161/01.STR.0000174487.81126.ef

Wischnewski, M., & Schutter, D. J. L. G. (2015). Efficacy and time course of theta burst stimulation in healthy humans. Brain Stimulation, 8(4), 685–692. https://doi.org/10.1016/J.BRS.2015.03.004

